## Supplementary Information for "Neural circuits in the mouse retina support color vision in the upper visual field"

Szatko, Korympidou et al.

- Supplementary Statistical Analysis
- Supplementary Figures
- Supplementary References

### SUPPLEMENTARY STATISTICAL ANALYSIS

#### Linear Mixed-Effects Models

We used a Linear Mixed-Effects Model to analyze the difference between center and surround spectral contrast (*SC)*. This allowed to incorporate a random effect term accounting for the fact that not all ROIs with a center response displayed a surround response (partially paired data).

$$y = X\beta + Zu + \epsilon$$

Here, *y* is the dependent variable *SC*, $\beta$ is the coefficient vector for the fixed effects (e.g. center/surround and retinal location), *u* is the coefficient vector of random effects (here: cell ID), $\epsilon$ is an unknown vector of random errors and *X* and *Z* are known design matrices relating the observations *y* to $\beta$ and $\epsilon$, respectively.

We used the `lmerTest`-package (version 3.0.1) for R to implement the model and perform statistical testing (Bates et al., 2015).

<u>Center and surround *SC* for OPL recordings</u>

For cones in OPL recordings (Fig. 2b, c), we modeled *SC* as a function of center and surround ("cent_surr") and cell ID.

```
SC ~ cent_surr + (1|cell_id)
```

For dorsal scan fields, the resulting model was fit using n=922 observations, with n=689 paired observations in group cell_id. Running an ANOVA on the model yielded the following results:

```
Type III Analysis of Variance Table with Satterthwaite's method
           Sum Sq   Mean Sq   NumDF   DenDF   F value   Pr(>F)
cent_surr  377.62   377.62    1       920     3.7622    0.05273
```

The P value ~0.05 (effect size=0.37, s.d. error=0.38) indicates that for dorsal cones center *SC* is at the threshold of significance.

For ventral scan fields, the resulting model was fit using n=2,181 observations, with n=1,344 paired observations in group cell_id. Running an ANOVA on the model yielded the following results:

```
Type III Analysis of Variance Table with Satterthwaite's method
           Sum Sq   Mean Sq   NumDF   DenDF   F value   Pr(>F)
cent_surr  1730.5   1730.5    1       2179    178.8     2.2e-16
```

The model results indicate that for ventral cones center *SC* is significantly different from surround *SC* (effect size=-0.63, s.d. error=0.08).

<u>Center and surround *SC* for IPL and GCL recordings</u>

For ROIs in IPL (Fig. 4b) and GCL (Fig. 6b) recordings, we modeled *SC* with factors cent_surr and retinal location ("bin_num") as well as their interaction and cell ID as random effect.

```
SC ~ cent_surr * ret_loc + (1|cell_id)
```

For IPL ROIs, the resulting model was fit using n=6,143 observations, with n=3,188 observations in group cell ID. Running an ANOVA on the model yielded the following results:

```
Type III Analysis of Variance Table with Satterthwaite's method
                Sum Sq  Mean Sq   NumDF   DenDF    F value    Pr(>F)
cent_surr       384.14  384.14    1       6139     2863.77    <2e-16
bin_number      23.68   23.68     1       6139     2863.77    <2e-16
cent_surr:bin   264.73  264.73    1       6139     2863.77    <2e-16
```

The model results indicate that for IPL ROIs there is a significant interaction between center and surround and retinal location (as is evident from Fig. 4b). Post-hoc testing using the `lsmeans`-package (version 2.30.0) for R revealed that center and surround *SC* are significantly different across all retinal locations:

```
bin_num_str =   one:
  contrast            estimate   SE         df    t.ratio    p.value
  center - surround   -0.660     0.0147     3094   -45.018   <.0001
bin_num_str =   two:
  contrast            estimate   SE          df   t.ratio    p.value
  center - surround   -0.830     0.0343     3073   -24.225   <.0001
bin_num_str =   three:
  contrast            estimate   SE         df    t.ratio    p.value
  center - surround   -0.657     0.0288     3154  -22.801    <.0001
bin_num_str =   five:
  contrast            estimate   SE         df    t.ratio    p.value
  center - surround   -0.542     0.0280     3248  -19.367    <.0001
bin_num_str =   six:
  contrast            estimate   SE         df    t.ratio    p.value
  center - surround   0.231      0.0211     3221  10.912     <.0001
bin_num_str =   seven:
  contrast            estimate   SE         df    t.ratio    p.value
  center - surround   0.347      0.0294     3048  11.788     <.0001
bin_num_str =   eight:
  contrast            estimate   SE         df    t.ratio    p.value
  center - surround   0.258      0.0228     3026  11.293     <.0001
```

For GCL ROIs, the resulting model was fit using n=10,804 observations, with n=5,788 observations in group cell ID. Running an ANOVA on the model yielded the following results:

```
Type III Analysis of Variance Table with Satterthwaite's method
                Sum Sq  Mean Sq   NumDF   DenDF    F value    Pr(>F)
cent_surr       441.82  441.82    1       10800    1375.51    <2e-16
bin_number      33.58   33.58     1       10800    104.53     <2e-16
cent_surr:bin   77.65   77.65     1       10800    241.73     <2e-16
```

The model results indicate that for GCL ROIs there is a significant interaction between center and surround and retinal location (as is evident from Fig. 6b). Post-hoc testing revealed that center and surround *SC* are significantly different across all retinal locations:

```
bin_num_str =   one:
  contrast            estimate   SE          df   t.ratio    p.value
  center - surround   -0.6824    0.0456      5558 -14.969    <.0001
bin_num_str =   two:
  contrast            estimate   SE          df   t.ratio    p.value
  center - surround   -0.6281    0.0218      5525 -28.768    <.0001
bin_num_str =   three:
  contrast            estimate   SE          df   t.ratio    p.value
  center - surround   -0.4851    0.0188      5473 -25.853    <.0001
bin_num_str =   four:
  contrast            estimate   SE          df   t.ratio    p.value
  center - surround   -0.5655    0.0383      5626 -14.762    <.0001
bin_num_str =   five:
  contrast            estimate   SE          df   t.ratio    p.value
  center - surround   0-0.9121   0.0615      5972 -14.827    <.0001
bin_num_str =   six:
  contrast            estimate   SE          df   t.ratio    p.value
  center - surround   -0.2191     0.0291     5903 -7.535     <.0001
bin_num_str =   seven:
  contrast            estimate   SE          df   t.ratio    p.value
  center - surround   -0.0653    0.0332      5631 -1.966     <.0001
```

### Generalized Additive Models

We used Generalized Additive Models (GAMs) to analyze the relationship of difference in center and surround *SC* ($SC_{Diff}$) and IPL depth; opponency and IPL depth; center *SC* ($SC_{center}$) and IPL depth. GAMs extend the generalized linear model by allowing the linear predictors to depend on arbitrary smooth functions of the underlying variables (Wood, 2006):

$$g(\mu) = \beta_0 + f_1(x_1) + \ldots + f_n(x_n)$$

Here, $x_i$ are the predictor variables, $g$ is a link function and the $f_i$ are smooth functions of the predictor variables.

In practise, we used the `mgcv`-package for R (version 1.8-24) to implement GAMs and perform statistical testing.

<u>$SC_{Diff}$ vs. IPL depth</u>

To model the dependence of $SC_{Diff}$ as a function of IPL depth for dorsal and ventral IPL ROIs (Fig. 4e), we used a Gaussian GAM with factor retinal position ("dv"; 0: ventral; 1: dorsal) and a smooth term for dorsal and ventral retina as a function of IPL depth.

```
SC_Diff ~ s(Depth, by = dv, k = 20) + dv + s(exp_num, bs = "re")
```

We set the basis dimension k=20 to allow for sufficiently "wiggly" smooth terms. Inspection of the model fit indicated that this was high enough (using gam.check). The resulting model was fit using n=2,955 data points and yielded the following results:

```
Parametric coefficients:
                 Estimate   Std. Error  t value   Pr(>|t|)
(Intercept)      -0.04636   0.01391     -3.33     0.000871
dvV               0.98749   0.01696     58.214    < 2e-16
Approximate significance of smooth terms:
                 edf        Ref. df     F         p-value
s(Depth):dvD     1.0000     1.00        13.45     0.000249
s(Depth):dvV     11.5912    14.04       23.83     < 2e-16
```

Thus, the smooth terms for dorsal and ventral ROIs are highly significant, indicating non-random variation of $SC_{Diff}$ with IPL depth.

Overall, the model explained 57.2% of the deviance.

<u>Opponency vs. IPL depth</u>

To model the dependence of opponency as a function of IPL depth for dorsal and ventral IPL ROIs (Fig. 4f), we used a binomial GAM with factor retinal position ("dv"; 0: ventral; 1: dorsal) and a smooth term for dorsal and ventral retina as a function of IPL depth.

```
Opponency ~ s(Depth, by = dv, k = 20) + dv + s(exp_num, bs = "re")
```

The resulting model was fit using n=3,188 data points and yielded the following results:

Parametric coefficients:

| | Estimate | Std. Error | z value | Pr(>\|z\|) |
|---|---|---|---|---|
| (Intercept) | -3.1234 | 0.1745 | -17.9 | < 2e-16 |
| dvV | 3.4335 | 0.1826 | 18.8 | < 2e-16 |

Approximate significance of smooth terms:

| | edf | Ref. df | Chi. sq | p-value |
|---|---|---|---|---|
| s(Depth):dvD | 6.0127 | 7.500 | 38.88 | 3.66e-06 |
| s(Depth):dvV | 5.9131 | 7.419 | 211.14 | < 2e-16 |

The smooth terms for dorsal and ventral ROIs are highly significant, indicating non-random variation of number of opponent ROIs with IPL depth.

Overall, the model explained 30.8% of the deviance.

$SC_{center}$ vs. IPL depth

To model the dependence of $SC_{center}$ as a function of IPL depth for dorsal and ventral IPL ROIs (Suppl. Fig. S2b), we used a Gaussian GAM with factor retinal position ("dv"; 0: ventral; 1: dorsal) and a smooth term for dorsal and ventral retina as a function of IPL depth.

$SC_{center}$ ~ s(Depth, by = dv, k = 20) + dv + s(exp_num, bs = "re")

The resulting model was fit using n=3,188 data points and yielded the following results:

Parametric coefficients:

| | Estimate | Std. Error | z value | Pr(>\|z\|) |
|---|---|---|---|---|
| (Intercept) | 0.1837 | 0.0046 | 39.68 | < 2e-16 |
| dvV | -0.6771 | 0.0057 | -118.22 | < 2e-16 |

Approximate significance of smooth terms:

| | edf | Ref. df | Chi. sq | p-value |
|---|---|---|---|---|
| s(Depth):dvD | 9.1231 | 11.23 | 4.078 | 4.85e-06 |
| s(Depth):dvV | 11.1562 | 13.57 | 11.092 | < 2e-16 |

The smooth terms for dorsal and ventral ROIs are highly significant, indicating non-random variation of $SC_{center}$ with IPL depth.

Overall, the model explained 82% of the deviance.

### SUPPLEMENTARY FIGURES

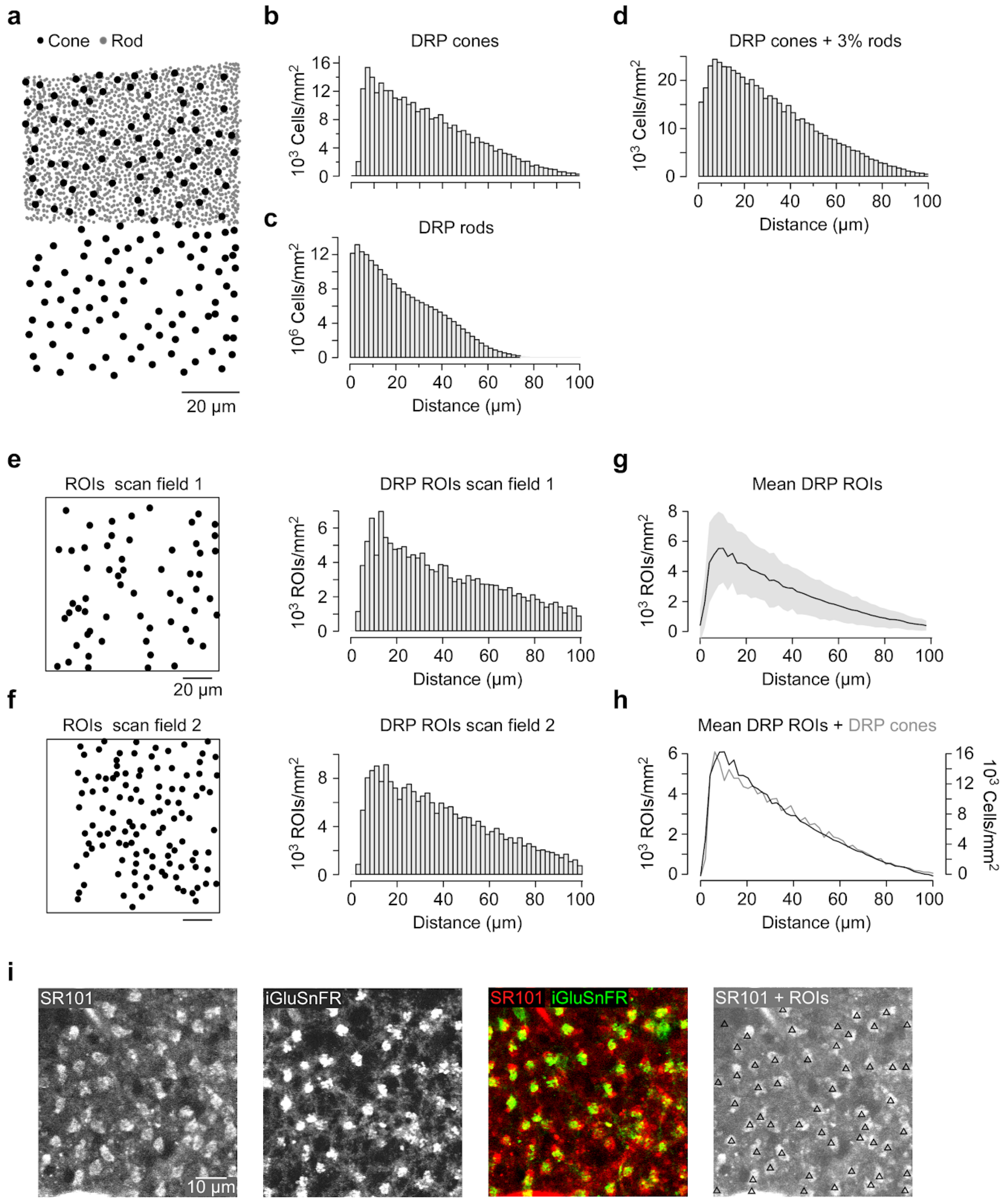


**Suppl. Figure S1 | Contribution of rod photoreceptors to glutamate signals in the outer plexiform layer. a,** Center-of-mass of cone (black) and rod (grey) axon terminals from electron-microscopy reconstruction (Behrens et al., 2016). Note that rod axon terminals were only reconstructed for the upper half of the retinal patch. **b, c,** Density recovery profile (DRP; (Rodieck, 1991)) for cones (b) and rods (c) shown in (a), with 2 µm bins. Note the low number of cells for distances <5 µm for the cone DRP, indicating a regular mosaic array. **d,** DRP of cones from (a) with 3% of rods randomly distributed across the retinal patch. Note that including a low percentage of rods reduces the regularity of the mosaic, indicated by neighboring cells with distances <5 µm. **e,** Exemplary scan field with ROI positions indicated in black and corresponding DRP (right). **f,** Like (e) for a different scan field. **g,** Mean DRP of all OPL scan fields (n=52 scan fields, n=9 mice) with s.d. shading in grey. **h,** DRP from anatomical cone data from (b; grey) and mean DRP from (g; black) are not significantly different. p>0.05; Chi-squared test. **i,**

Sulforhodamine 101 (SR101) labeling of cone axon terminals (left; (Chapot et al., 2017)), iGluSnFR labeling (middle left), overlay image (middle right) and ROI position (black triangles) on top of SR101 labeling (right) for an exemplary scan field. The right image illustrates that functional glutamate release units (ROIs) match anatomically identified cone axon terminals (SR101).

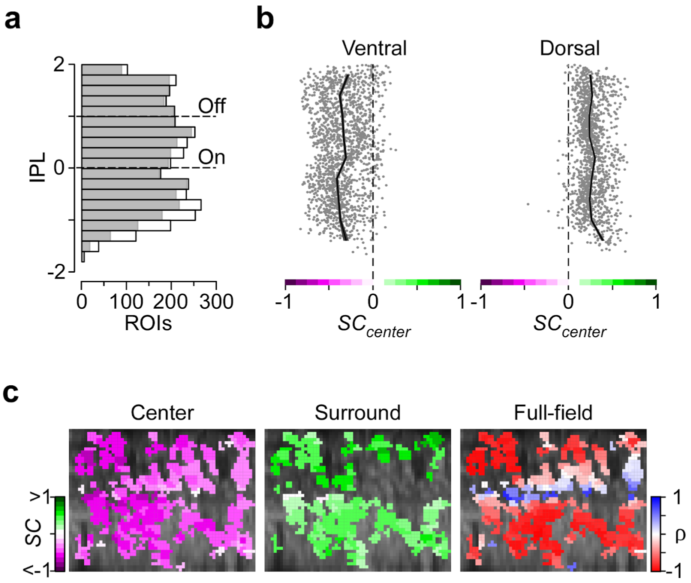


**Suppl. Fig. S2 | Chromatic glutamate responses across the inner plexiform layer. a,** Distribution of all recorded ROIs across the IPL (black), with ROIs that passed our quality criterion indicated in grey (for details, see Methods) and On and Off ChAT bands indicated by dashed line. Bin size: 0.4. **b,** Receptive field (RF) center spectral contrast ($SC_{center}$) values across the IPL for ROIs located in the ventral (left) and dorsal (right) retina, respectively. For statistics, see Suppl. Information. **c,** Example scan field located in the ventral retina, with ROIs color-coded according to RF center (left) and surround *SC* (middle) as well as linear correlation coefficient (ρ) of full-field events (right). For most scan fields in the ventral retina, the difference between center and surround *SC* and the fraction of color-opponent responses was more pronounced in the IPL's Off sublamina. However, for a small number of ventral scan fields, this was not the case.

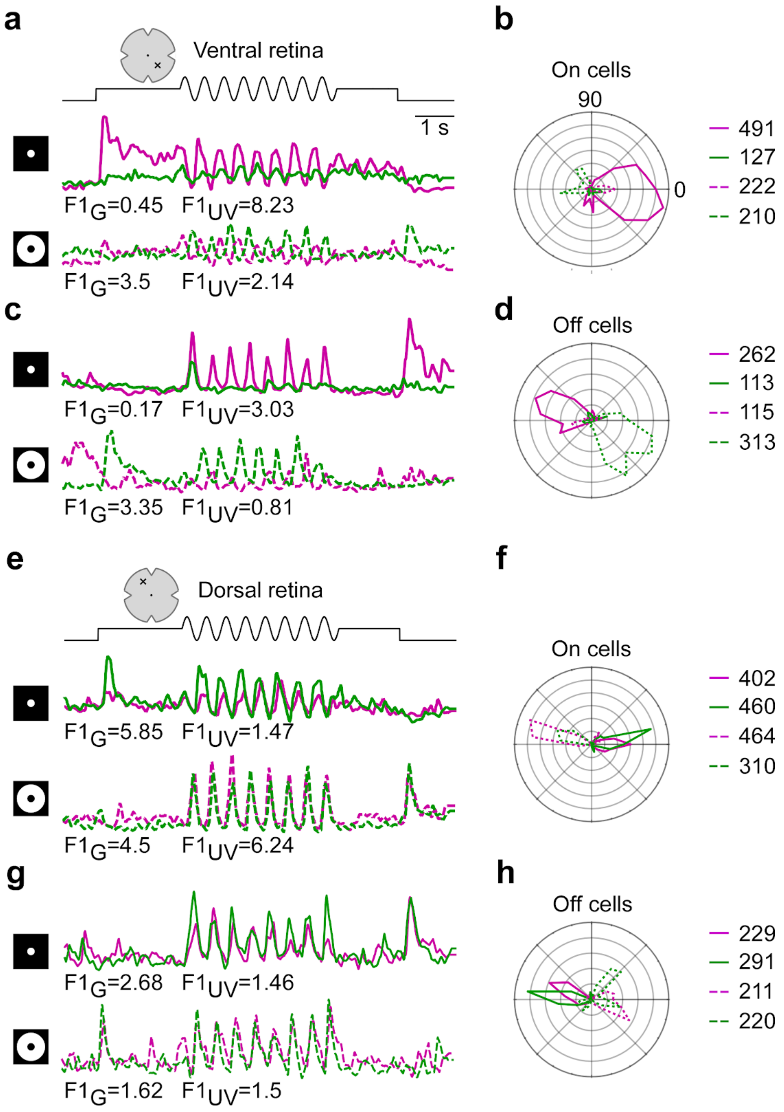


**Suppl. Fig. S3 | Bipolar cell responses to sinusoidal modulation of green and UV LED. a,** Mean glutamate trace (n=3 trials) of an exemplary ROI located in the On layer of the IPL in the ventral retina in response to 2 Hz center (top) and surround (bottom, dashed lines) modulation of green and UV LED. The fundamental response component (F1) is indicated below the traces. **b,** Polar plot showing distribution of response phases (in degrees) for On cells located in the ventral retina to stimulus shown in (a). Each histogram is normalized according to mean F1 (for details, see Methods). Numbers on the right indicate ROIs used for analysis of each condition. **c,d,** Like (a,b), but for Off cells located in the ventral retina. **e,f,** Like (a,b), but for On cells located in the dorsal retina. **g,h,** Like (a,b), but for Off cells located in the dorsal retina.

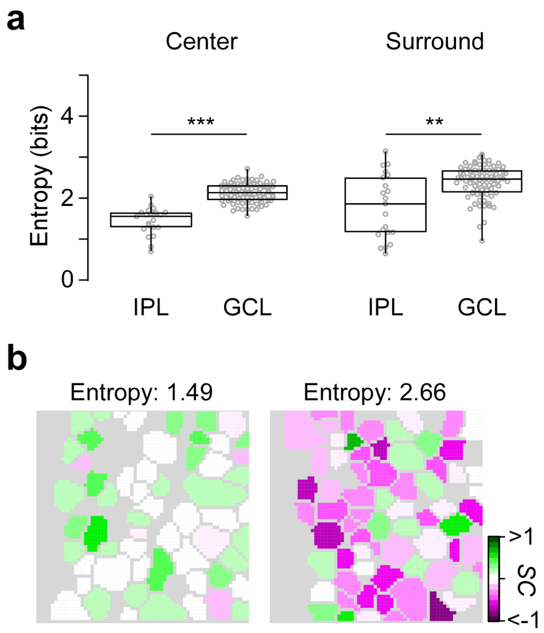


**Suppl. Fig. S4 | Diversity of chromatic preferences within IPL and GCL scan fields. a,** Distribution of field entropy of IPL (n=21) and GCL (n=82) scan fields using center (left) and surround (right) spectral contrast. High field-entropy indicates high chromatic tuning heterogeneity within single scan fields. **: $p<0.01$; ***: $p<0.001$; Wilcoxon signed-rank test. **b,** Two exemplary GCL scan fields with low (left) and high (right) entropy.

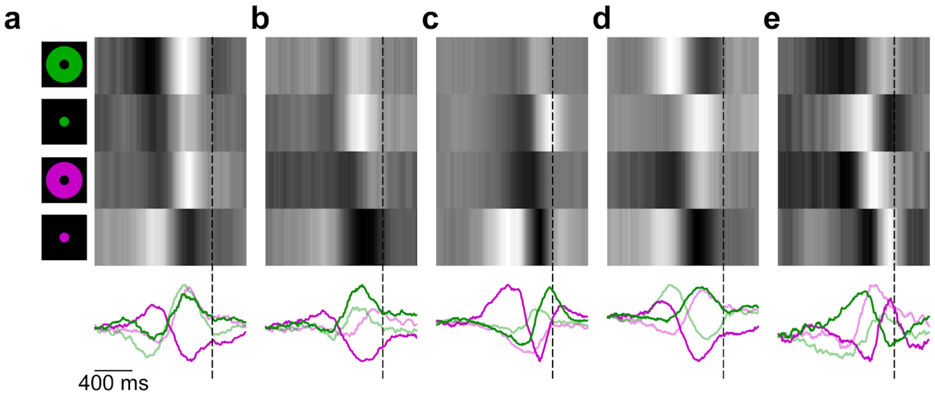


**Suppl. Fig. S5 | Center-opponent cells in the ganglion cell layer of the mouse. a,** Response-triggered stimulus kernels for UV and green center and surround stimulation (top) of exemplary center-opponent GCL cells, with color-coded center (bright) and surround (dim) kernels overlain (bottom). Dashed line corresponds to time point of response. **b-e,** Like (a), but for other center-opponent GCL cells. Note the differences in response polarity and kinetics, suggesting that center-opponent GCL cells do not comprise a single functional type.

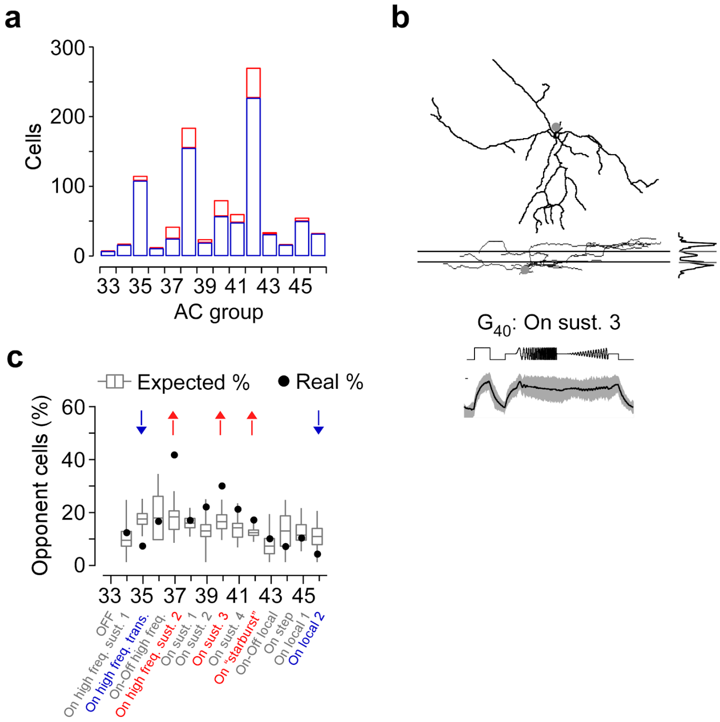


**Suppl. Fig. S6 | Center-opponent displaced amacrine cells (dAC). a,** Distribution of color-opponent (red) and non-opponent (blue) dACs located in the ventral retina. For further analysis, only groups with n>10 cells were used. **b,** Dendritic morphologies with stratification profiles of exemplary color-opponent cell assigned to dAC group $G_{40}$, dye-filled and reconstructed subsequent to imaging experiments. Lines for side-view of morphology and stratification profile indicate On and Off ChAT bands. Mean full-field chirp responses (black, s.d. shading in grey) of the same groups indicated below. **c,** Box plots show distribution of expected percentages of color-opponent cells given center *SC* and $SC_{Diff}$ values in each group (for details, see Methods). Black circles indicate true percentage of color-opponent cells. Arrows pointing down- and up indicate groups with significantly more and less color-opponent cells than expected, respectively.

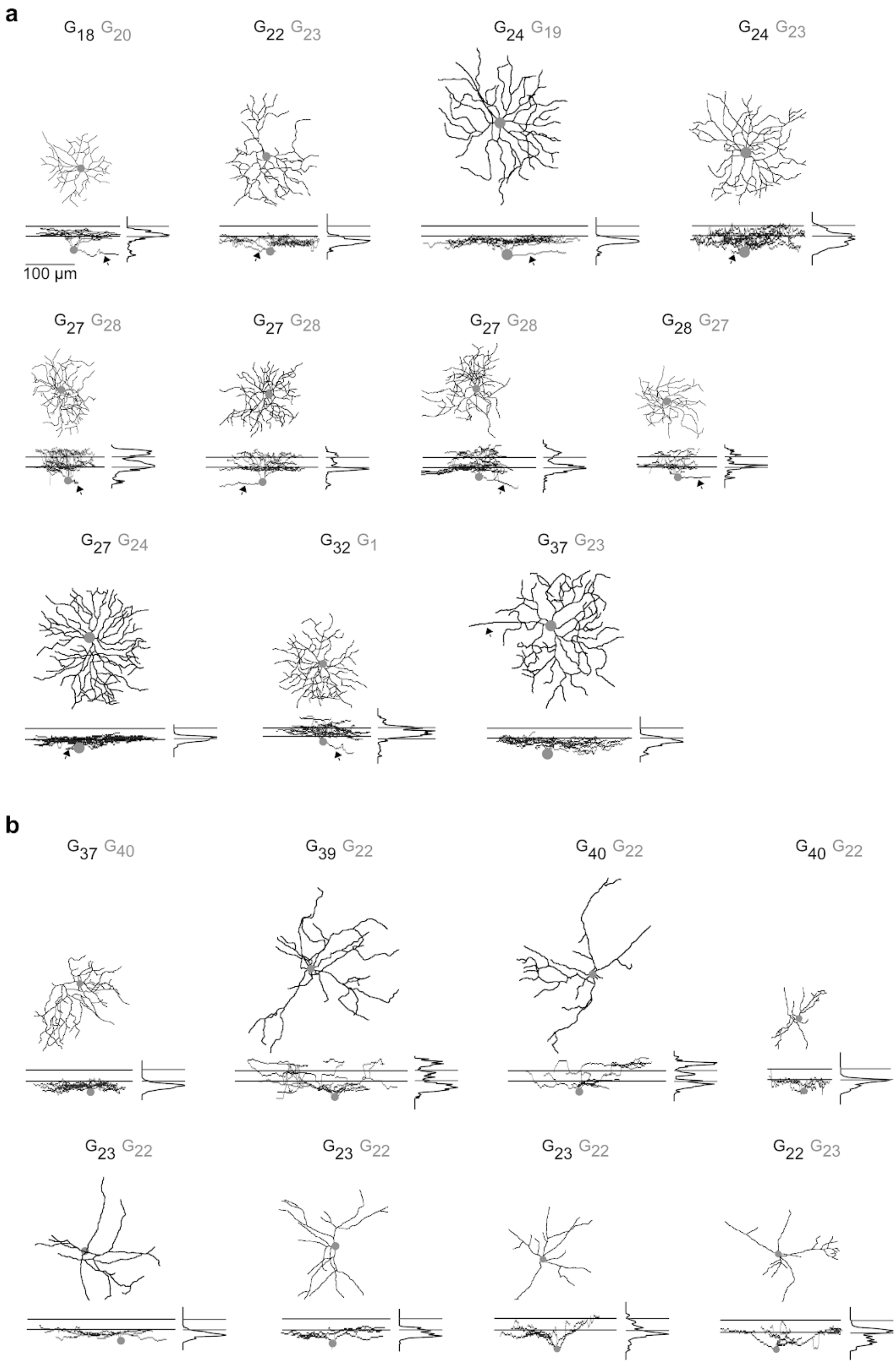


**Suppl. Fig. S7 | Morphologies of color-opponent GCL cells**. **a,** Top-view and side-view of dye-injected and reconstructed color-opponent RGCs identified by the presence of an axon (arrow), with most (black) and second-most (grey) likely group assignment indicated above morphologies. **b,** Same as (a), but for dACs.

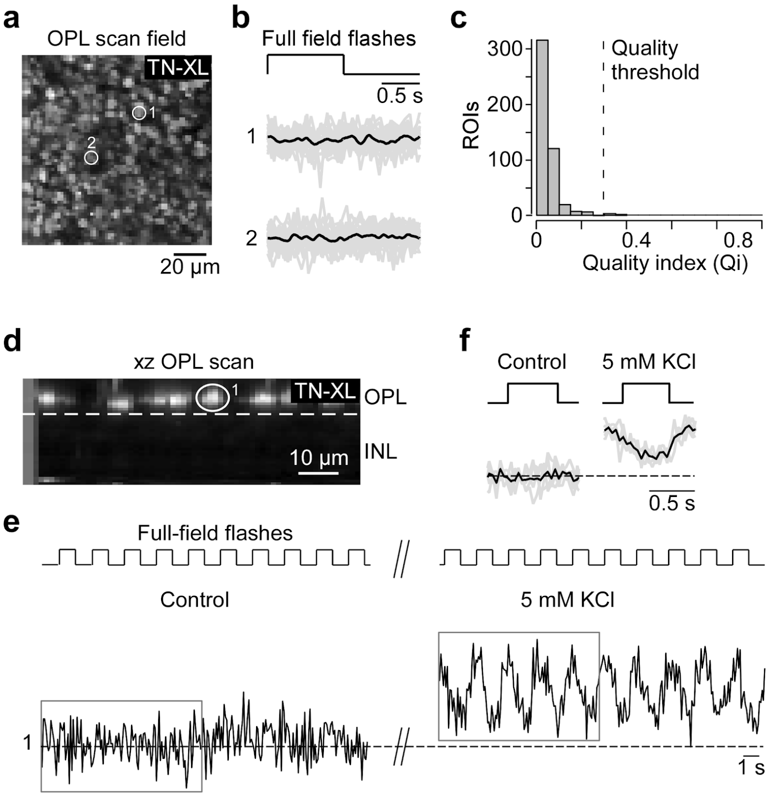


**Suppl. Figure S8 | Calcium imaging in the OPL using the biosensor TN-XL. a,** Example scan field of cone axon terminals expressing the calcium indicator TN-XL in the outer plexiform layer (OPL) of a whole-mounted HR2.1:TN-XL mouse retina (Wei et al., 2012), with two axon terminals indicated (circles). **b,** Mean calcium responses (black) with individual trials (grey) to full-field flashes (700 µm in diameter) of the cones indicated in (a). **c,** Histogram of quality indices (Qi) of full-field flash responses (n=370 ROIs, n=4 scan fields, n=4 mice). In contrast to recordings in retinal slices (Baden et al., 2013; Kemmler et al., 2014; Chapot et al., 2017), we did not detect any light-evoked calcium changes in cone axon terminals. Dashed line indicates Qi threshold used for analysis of iGluSnFR data (cf. Fig. 2). **d,** Example scan field of a vertical optical slice recording using an electrically tunable lens (ETL), with one axon terminal indicated. Dotted line indicates OPL border. **e,** Raw calcium trace of cone shown in (d) in response to full-field flashes for control condition (left; 2.5 mM KCl) and with increased KCl concentration (right; 5 mM KCl). Increasing the extracellular potassium concentration, which is expected to slightly depolarize the cones, recovered their light responses. This suggests that the lack of robust light responses is likely due to the limited dynamic range of TN-XL (discussed in (Wei et al., 2012)). The difference to the results in slices is probably due to increased laser-evoked activity in the retinal whole-mount preparation (discussed in (Euler et al., 2009, 2019)). Dashed line indicates calcium baseline of control condition. Grey rectangles indicate time used for averages shown in (f). **f,** Mean calcium trace (black) with individual trials (grey) for control and 5 mM KCl.

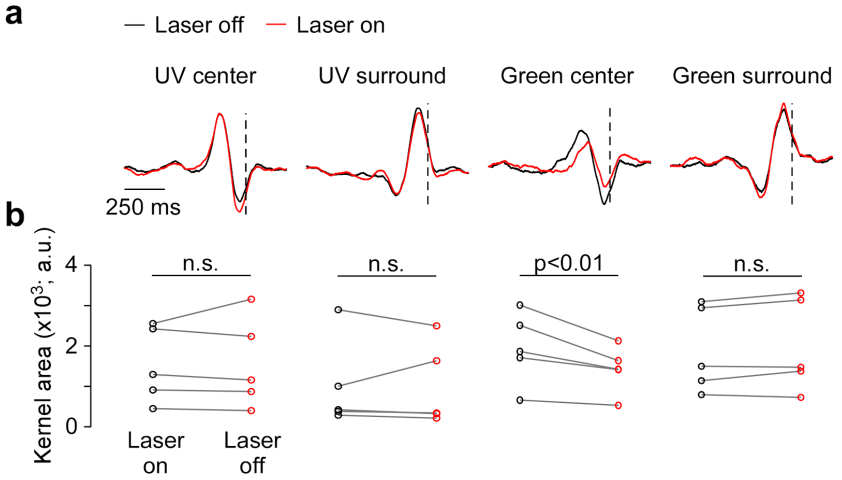


**Suppl. Fig. S9 | Effect of the two-photon laser on chromatic RGC responses. a,** Spike-triggered response kernels for UV and green center and surround stimulation of an exemplary RGC electrically recorded without (black) and with (red) simultaneously scanning the tissue with the two-photon laser. For detailed discussion of laser-induced activity in the retina, see (Euler et al., 2009, 2019). **b,** Areas of UV and green center and surround kernels for n=12 electrically recorded RGCs. The two-photon laser consistently reduced the green center component of RGC RFs by ~25%, while it had no detectable effect on UV center or UV and green surround kernels. These results indicate that green-sensitive visual pigments are more strongly activated by the laser than UV-sensitive ones. The effect is restricted to the RF center probably because of the relatively small size of the recording fields. This effect results in an underestimation of the green center component of RFs in our dataset but it does not change the conclusions of this study.
